# Breaking the fibrinolytic speed limit with microwheel co-delivery of tissue plasminogen activator and plasminogen

**DOI:** 10.1101/2021.05.05.440940

**Authors:** Dante Disharoon, Brian G. Trewyn, Paco S. Herson, David W.M. Marr, Keith B. Neeves

## Abstract

Fibrinolysis is the enzymatic degradation of fibrin, the biopolymer that gives blood clots their mechanical integrity. To reestablish blood flow in vessels occluded by clots, tissue plasminogen activator (tPA) can be used; however, its efficacy is limited by transport to and into a clot and by the depletion of its substrate, plasminogen. To overcome these rate limitations, we design a platform to co-deliver tPA and plasminogen based on microwheels (μwheels), wheel-like assemblies of superparamagnetic colloidal beads that roll along surfaces at high speeds and carry therapeutic payloads in applied magnetic fields. By experimentally measuring fibrinolysis of plasma clots at varying concentrations of tPA and plasminogen, the biochemical speed limit was first determined. These data, in conjunction with measurements of μwheel translation, activity of immobilized tPA on beads, and plasminogen release kinetics from magnetic mesoporous silica nanoparticles (mMSN), were used in a mathematical model to identify the optimal tPA:plasminogen ratio and guide the coupling of plasminogen-loaded mMSN to tPA functionalized superparamagnetic beads. Once coupled, particle-bead assemblies form into a co-delivery vehicle that rolls to plasma clot interfaces and lyses them at rates comparable to the biochemical speed limit. With the addition of mechanical action provided by rotating μwheels to penetrate clots, this barrier was exceeded by rates 40-fold higher lysis by 50 nM tPA. This co-delivery of an immobilized enzyme and its substrate via a microbot capable of mechanical work has the potential to target and rapidly lyse clots that are inaccessible by mechanical thrombectomy devices or recalcitrant to systemic tPA delivery.

## Introduction

Microbots are promising therapeutic vehicles because of their potential to target drug delivery to disease-afflicted sites within the body.^1^ Their propulsion can be achieved using applied magnetic,^2,3^ electric,^4^ optic,^5^ and acoustic fields.^6,7^. Of these strategies, magnetic field based propulsion approaches are particularly well suited for in vivo applications because magnetic fields do not attenuate in tissue^8^ and are not harmful to the human body. As a result, magnetically-controlled microbots have been proposed for the treatment of cancer,^9,10^ ocular surgery,^11^ and tissue damage^12^ where targeted drug delivery could provide substantial benefit. Building on these studies, we have recently shown that microbots can not only deliver drug but also impart mechanical action to aid in drug efficacy, specifically in the targeting of blood clots for the treatment of stroke.^13^

Fibrinolysis is the biochemical breakdown of fibrin, which is the biopolymer that gives clots their mechanical integrity.^14^ It can be used to reestablish blood flow in various types of thrombosis including myocardial infarction,^15^ deep vein thrombosis,^16^ pulmonary embolism,^17^ ischemic stroke,^18,19^ and limb ischemia.^20^ Recombinant tissue plasminogen activators (tPAs) are the only currently Food and Drug Administration approved thrombolytic drug. They work by binding to and converting the zymogen plasminogen to the enzyme plasmin, an interaction that is greatly accelerated in the presence of its co-factor fibrin.^21^ Plasmin in turn cleaves lysyl and arginyl bonds at many different sites to transect cross-linked fibrin fibers producing several different fragments.^22^

The rate of fibrinolysis by tPA is limited by both transport processes and biochemical reactions.^23^ The transport of tPA to the clot periphery can be the rate limiting step in intravenous delivery because it is diffusion-dominated in occluded vessels with low or no blood flow (Fig. 1A).^24,25^ tPA (alteplase) has a half-life of minutes in blood due to inactivation by its endogenous inhibitors plasminogen activator inhibitors 1 and 2 and thus is often eliminated before it reaches the thrombus.^26,27^ These limitations can be partially overcome by catheter directed intraarterial delivery, however only 20% of stroke patients have large artery occlusions that are accessible to catheters.^28^ Tenecteplase is an engineered tPA that is less susceptible to endogenous inhibitors and has a higher affinity for fibrin,^29^ but it has not shown superior performance to alteplase in clinical trials for ischemic stroke.^30^ Even once delivered, penetration of tPA is limited by its high binding affinity to fibrin fibers which localizes it to the first few micrometers of the thrombus interface.^31,32^ Acting as an effective affinity filter, this leads to surface erosion with little penetration of tPA into the clot. While a direct strategy of increasing tPA concentration to overcome slow diffusion rates would seem desirable, this is bounded by the bleeding risks associated with degradation of fibrinogen, plasminogen, and α2-antiplasmin, lysis of hemostatic clots, and tPA neurotoxicity.^33,34^

**Figure 1:**
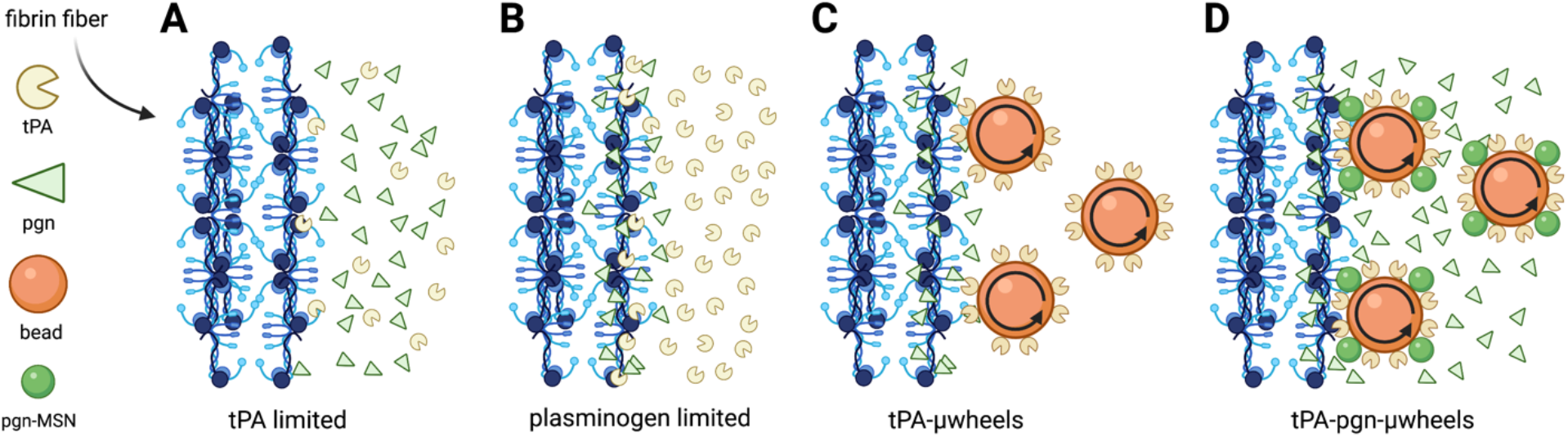
A) The rate-limiting steps of fibrinolysis include the transport to a clot and binding of tPA to fibrin fibers at low tPA concentrations. B) At sufficiently high tPA concentrations, its substrate plasminogen (pgn), becomes the limiting factor in fibrinolysis. C) These rate-limiting steps are overcome using magnetically powered μwheels, superparamagentic beads (orange spheres) that self-assemble in rotating magnetic fields. When μwheeels are coupled to tPA (tPA μwheels) they accumulate at the clot interface leading to high tPA concentrations and plasminogen limited fibrinolysis. D) By attaching plasminogen releasing nanoparticles (green spheres) to μwheels (tPA-pgn-μwheels) co-delivery of both enzyme and substrate are achieved yielding fibrinolysis rates that overcomes plasminogen-limited fibrinolysis.

Counterintuitively, the speed of fibrinolysis does not increase monotonically with tPA concentration. Rather, a bell-shaped curve has been reported with a maximum lysis rate occurring at ~20 nM tPA for compacted plasma clots.^35^ This is surprising because the plasma concentration of plasminogen (~2 μM) is much higher than the therapeutic tPA concentration (5-50 nM), an observation explained by consumption of *local* plasminogen due its high affinity binding to partially degraded fibrin and subsequent conversion to plasmin.^36–39^ Supplemental plasminogen and platelet derived plasminogen enhance the fibrinolysis rates^40,41^ suggesting that plasminogen is the limiting factor in the presence of high plasminogen activator concentrations (Fig. 1B).

The functionalization of micro and nanoparticles with tPA overcomes some of the transport and kinetic limitations of fibrinolysis by protecting tPA from inhibition and using targeting moieties to platelets and fibrin that localize particles on a clot while maintaining low circulating concentrations.^42^ Microparticles manipulated by external fields, or microbots, offer further advantages of not relying on blood flow for delivery.^43,44^ For example, tPA functionalized magnetic particles pulled towards a clot with a magnetic field gradient improves thrombolytic outcomes in mice^45^ and rats^46–48^ models of thrombosis compared to free tPA. Rotating magnetic microbots can also act as local mixers, reducing concentration gradients near the interface of a clot to accelerate tPA-mediated thrombolysis.^44^

In previous work with assemblies of superparamagnetic microparticles functionalized with tPA (tPA-μwheels), we have shown that actuation using rotating magnetic fields enables both delivery at high concentrations and mechanical disruption of clots, ultimately leading to an inside-out bulk erosion (Fig. 1C).^13^ However, while we were able to achieve local tPA concentrations that were three orders-of-magnitude higher than those by diffusive delivery of tPA, this resulted in only a one order-of-magnitude increase in fibrinolysis speed. Based on these results it appears that fibrinolysis using tPA-μwheels is plasminogen limited as reported in other studies of fibrinolysis at high tPA concentrations.^49^ To overcome this limitation and further enhance lysis speed, we develop here a μwheel-based strategy of delivering both tPA and plasminogen using tPA-μwheels coupled with mesoporous silica nanoparticles (MSN), which we have previously used to achieve controlled release of proteins.^50–52^ These co-laden μwheels break the biochemical speed limit by supplementing fresh plasminogen and burrowing their way into plasma-derived clots (Fig. 1D).

## Results

### Plasminogen depletion limits fibrinolysis rate at high tPA concentrations

To measure the extent to which fibrinolysis is rate-limited by tPA, we exposed a plasma clot formed with thrombin to varying concentrations of tPA and plasminogen (Fig. 2A). For a physiologic plasminogen concentration of 1 μM, the fibrinolysis rate, defined by the dissolution of the fibrin front over time, increases with increasing tPA concentrations from 50 nM up to 200 nM tPA (Fig. 2A, first column). However, at tPA concentrations > 200 nM the lysis rate decreases. Doubling the plasminogen concentration to 2 μM while holding tPA concentration at 50 nM resulted in a 65% increase in lysis rate and up to a 120% increase in lysis rate at tPA concentrations of 250-800 nM. Note that there is an optimal plasminogen concentration for a given tPA concentration that yields the maximum lysis rate (Fig. 2B). Plasminogen concentrations above this optimum attenuate fibrinolysis suggesting a competition for fibrin binding sites between tPA and plasminogen. Probing further, we measured fibrinolysis rates for a range of tPA:plasminogen ratios and identified accelerated fibrinolysis (AF) zones, defined as regions where lysis rates are within 10% of the maximum, or exceed 13.5 *μ*m/min (Fig. 2A). Here, it is apparent that there is a biochemical speed limit in the range to 200-600 nM tPA and 2-4 μM plasminogen, suggesting that a strategy of co-delivering tPA and plasminogen at these concentrations could significantly accelerate fibrinolysis. μWheels are well-suited for such a strategy because they are assembled from component, potentially-multifunctional, building blocks and can be driven to and accumulate at the edge of a clot, achieving a high local concentration of both the enzyme and its substrate adjacent to their co-factor.

**Figure 2:**
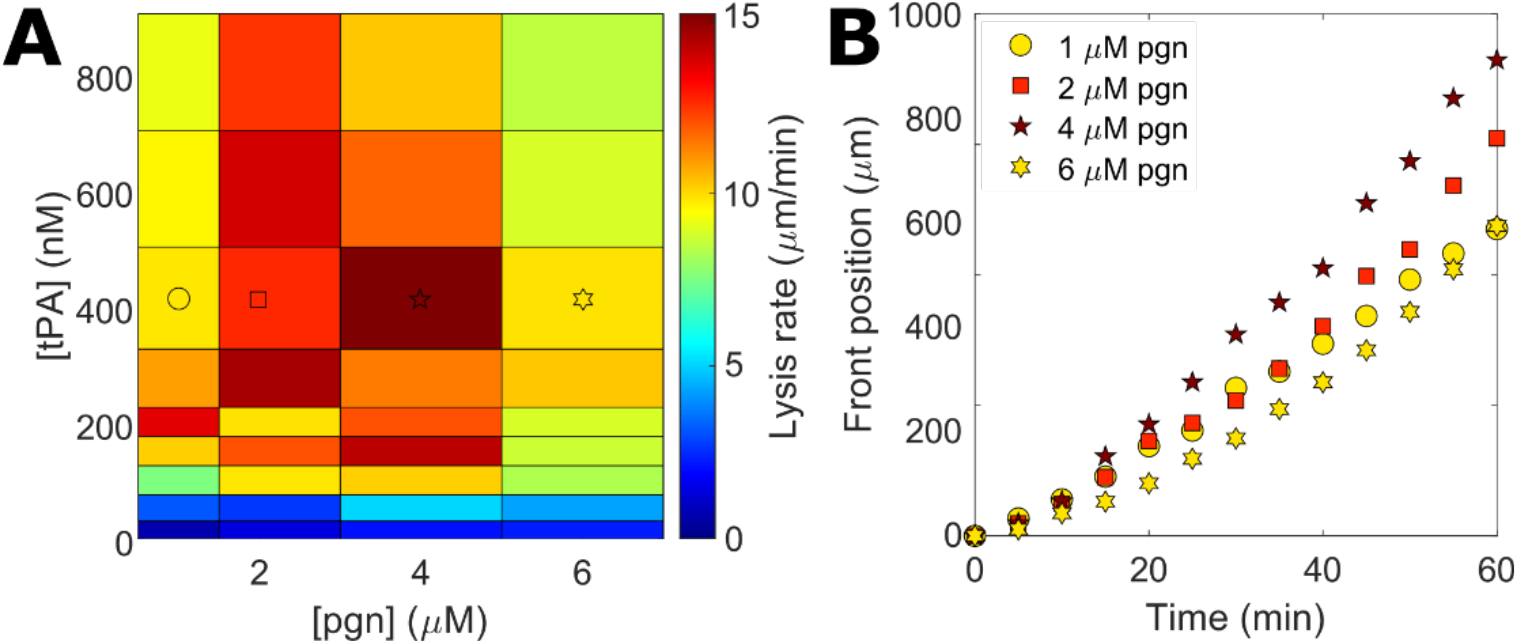
Measuring the fibrinolytic speed limit of plasma clots. A) Heat map of lysis rates as a function of tPA and plasminogen (pgn) concentrations. The bold border outlines the accelerated fibrinolysis (AF) zone. Symbols denote data shown in panel B. B) Dissolution of fibrin at 400 nM tPA with endogeneous plasma plasminogen concentration (1 μM) and the addition of exogenous plasminogen up to 6 μM total plasminogen concentration.

### Synthesis and characterization of plasminogen releasing magnetic colloids

To create tPA-μwheels, we functionalize streptavidin Dynabeads™ (beads) with biotinylated tPA (tPA-beads) (Fig. 3A). These tPA-beads readily assemble into tPA-μwheels with application of a magnetic field.^13^ To co-deliver plasminogen and tPA, we use mesoporous silica nanoparticles (MSN) with incorporated iron oxide nanoparticles to make them magnetic and increase coupling rate of beads and MSNs. We covalently coupled biotinylated magnetic mesoporous silica nanoparticles (mMSN) to beads to create studded beads, then functionalized the studded beads with tPA (tPA-studded beads) (Fig. 3A). Finally, the tPA-studded beads were incubated with plasminogen to load the mMSN to yield a superparamagnetic bead that can release plasminogen and carry immobilized tPA (pgn-tPA-beads) (Fig. 3A).

**Figure 3:**
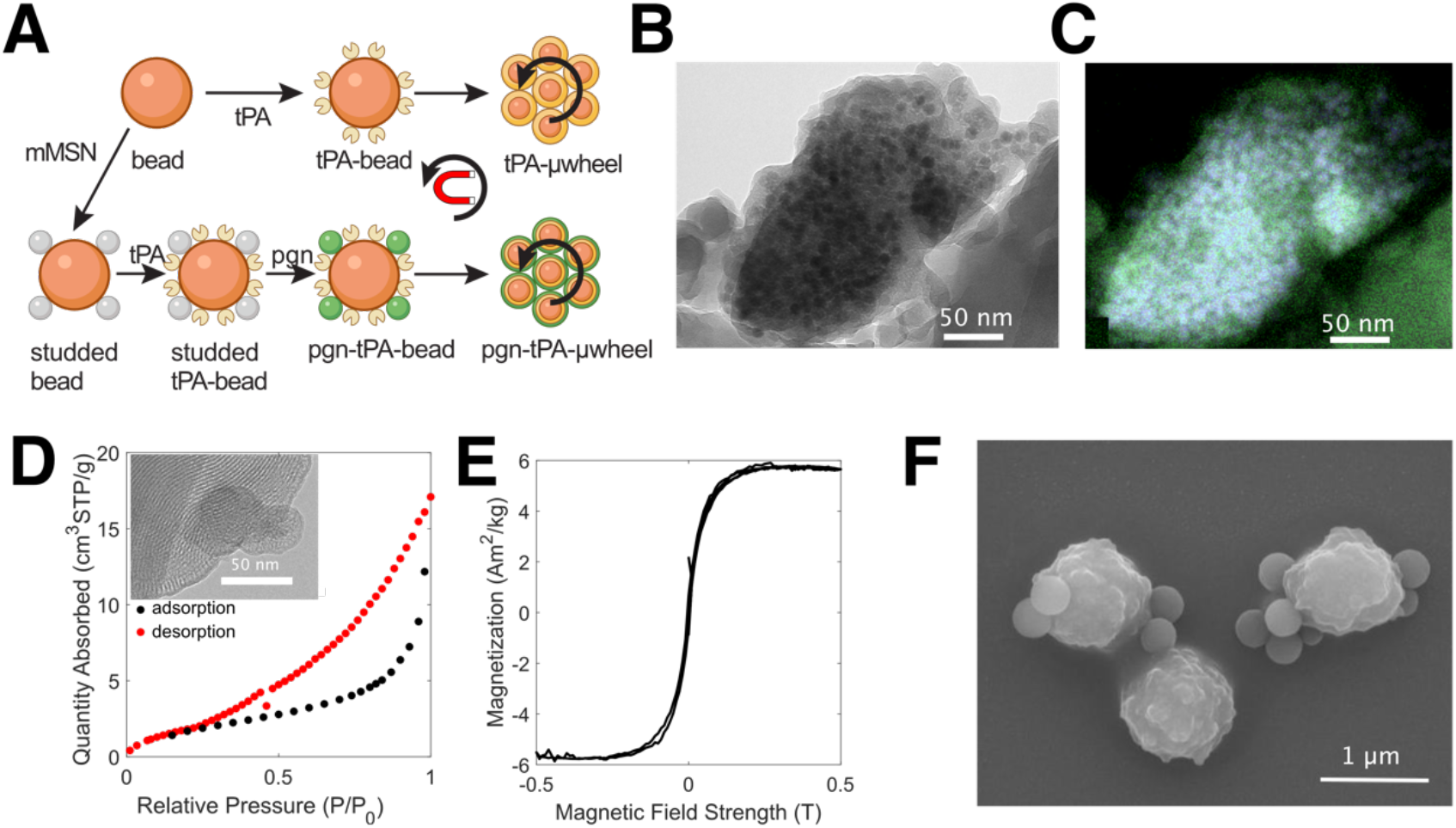
Synthesis and characterization of magnetic mesoporous silica nanoparticles (mMSN) and their coupling to superparamagnetic beads. A) Schematic of tPA-functionalization, mMSN bead coupling, and plasminogen (pgn) loading to create tPA and tPA-pgn-μwheels. B) Transmission electron microraph (TEM) of mMSN. Darker areas are Fe3O4 domains incorporated into the silica matrix. C) High-angle annular dark field energy-dispersive X-ray (HAADF-EDS) spectrum showing iron oxide domains (blue) distributed throughout silica matrix (green). D) Representative BJH isotherm data with TEM image of pore structure (inset). E) Magnetization profile for mMSN. F) Scanning electron micrograph (SEM) of studded beads.

The mMSN were synthesized with an average diameter 161 ± 57 nm and circularity 0.61 ± 0.17 and imaged using transmission electron microscopy (TEM) (Fig. 3B). Iron oxide content was measured at 10.5 wt% with 90% confidence using quantitative energy dispersive X-ray spectroscopy (EDS) (Fig. 3C). Nitrogen sorption isotherms show adsorption at relative pressures between 0.3 and 1.0 with most hysteresis occurring at 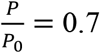 (Fig. 3D), indicating the presence of both micro and mesopores with a wide range of pore sizes. The Barrett-Joyner-Halenda (BJH) method was used to determine an accessible pore volume of 1.3 cm^3^/g distributed over a wide range of pore sizes with an average of 14.6 ± 5.7 nm. Magnetization curves for the mMSN confirmed that the particles are paramagnetic and amenable to magnetic control (Fig. 3E) allowing us to couple them to superparamagnetic beads with long-range magnetic attractions more efficiently than by mixing alone. Coupling between mMSN and beads was verified using scanning electron microscopy (SEM) (Fig. 3F).

### Designing co-delivery strategy for tPA and plasminogen

A strategy that relies solely on higher tPA concentrations does not necessarily lead to faster lysis (Fig. 2B). One of the features of the synthesis approach (Fig. 3A) is that the relative concentrations of tPA and plasminogen can be adjusted independently by varying the ratio of tPA-beads to plasminogen-loaded mMSN. To predict appropriate relative concentrations, we developed a mathematical model (see *Methods*) to estimate the local concentrations of tPA and plasminogen at a fibrin front. The model requires inputs of experimental data, presented below, to calculate μwheel number density at the clot interface and plasminogen release rate from mMSN.

Under the influence of rotating magnetic fields, both tPA-beads and pgn-tPA-beads self assemble into μwheels. The velocity profiles of μwheels consisting of various numbers of beads, or ‘-mers’, are shown in Fig. 4A-B. μWheels consisting of tPA-beads translate with a lognormal velocity profiles (Eqn 2, Fig. 4A) and an average velocity of 6.8 *μ*m/s. Larger tPA-*μ*wheels translate faster than smaller tPA-*μ*wheels as previously reported.^53^ μWheels consisting of pgn tPA-beads translate with a negatively skewed Gaussian velocity profile (Eqn 3, Fig. 4B) and an average velocity of 7.7 *μ*m/s. Differences here can be ascribed to asperities on the surface of beads that increase their translational velocity^54^ and the mMSNs on the pgn-tPA-beads likely perform in a similar fashion.

**Figure 4:**
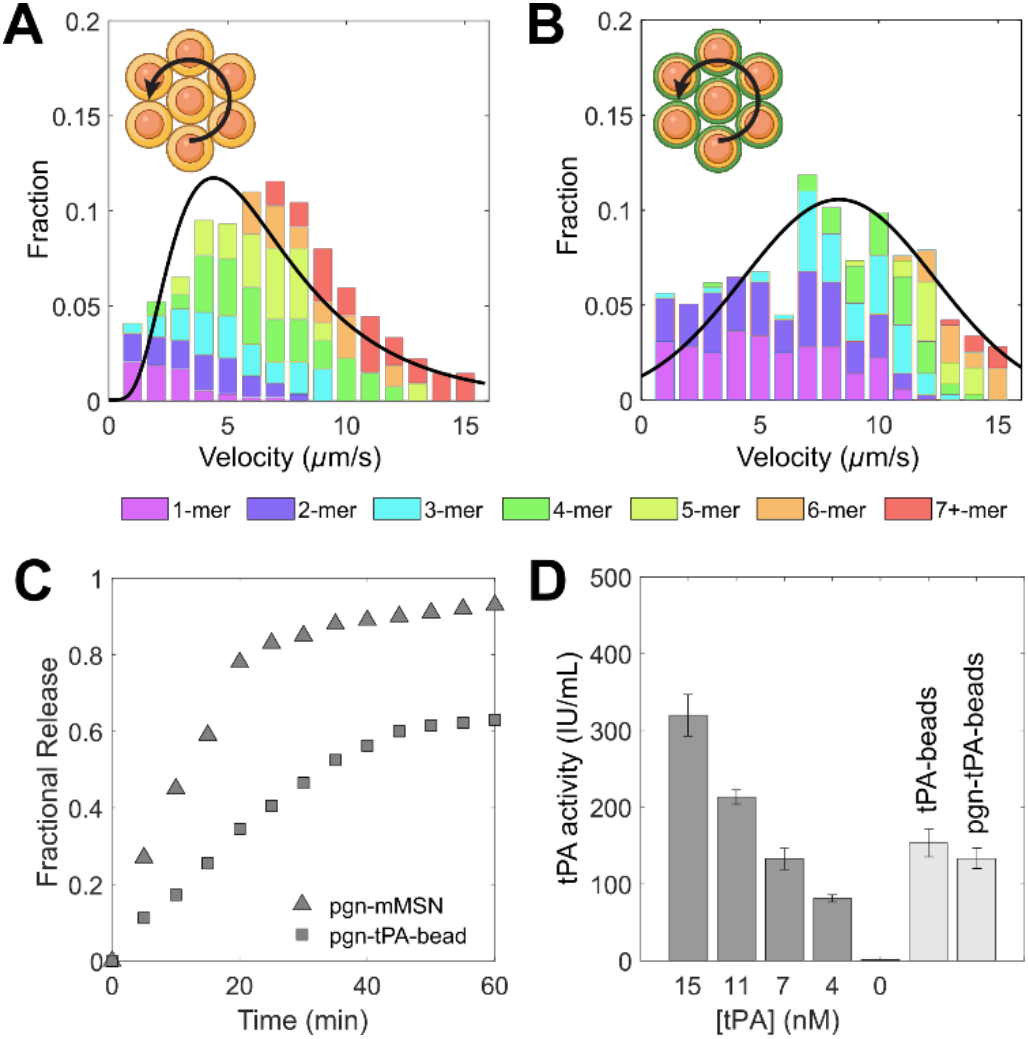
Representative velocity profile for tPA-beads (A) and pgn-tPA-beads (B). C) Plasminogen release profile for pgn-mMSN and pgn-tPA-beads at a number density of 10^6^/*μ*L. Fractional release is normalized to the total plasminogen loading as measured by spectrometry. D) Activity of 10^5^/μL bead populations compared to solvated tPA.

Plasminogen release kinetics from pgn-mMSN and pgn-tPA-beads were measured by UV absorbance (Fig. 4C). Pgn-tPA-beads have a lower loading capacity than pgn-mMSN, likely because some pores are inaccessible when coupled to beads. The amount of plasminogen released from pgn-tPA-beads was 66 ± 2% of that released from pgn-mMSN for a fixed mMSN number density. The activity of tPA and pgn-tPA-beads is equivalent to 8 nM tPA, a value not significantly affected by co-functionalization with pgn-mMSN (Fig. 4D). We measured the tPA activity on pgn-tPA-beads using a fluorogenic substrate as a function of coverage. As more mMSNs are bound to the beads, the surface available for tPA functionalization decreases as reflected in the decrease of tPA activity with higher coverage (Fig. 5A).

**Figure 5:**
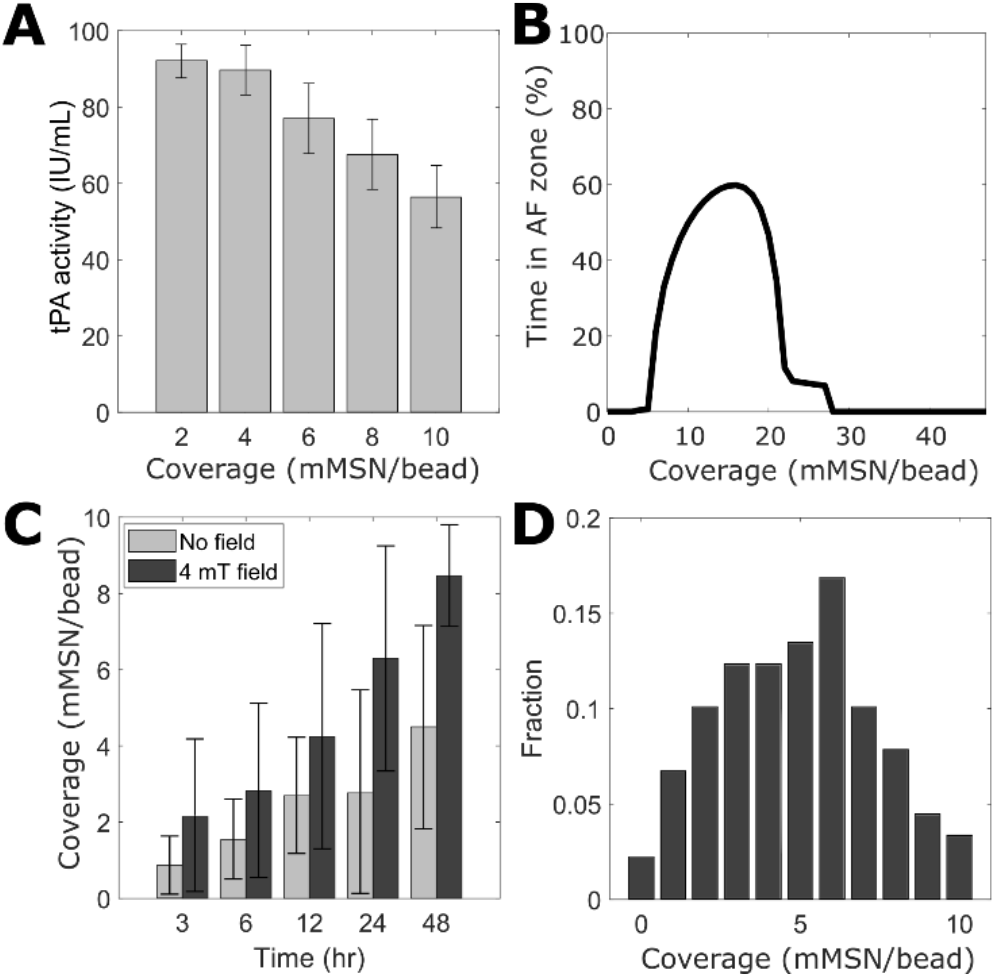
A) tPA activity as a function of coverage as measured using a fluorogenic substrate in a plate reader. B) Percentage of a 60 min experiment during which plasminogen concentration is within the accelerated fibrinolysis (AF) zones, defined as regions where lysis exceeds 13.5 *μ*m/min (see Fig. 2B). C) Coverage of mMSN to beads as a function of mixing time in the with and without a 4 mT magnetic field. D) Histogram of coverage for t = 24 hr in the presence of a 4 mT magnetic field (n=50). Data in (A) and (C) expressed as average and standard deviation of n = 3 and 25, respectively.

Mass action kinetics were used to model the binding of tPA and plasminogen to fibrin, as well as the conversion of plasminogen to plasmin (Eqns 6-9). The velocity (Fig. 4A), plasminogen release (Fig. 4C), and tPA activity (Figs 4D, 5A) data are used to calculate the rate of transport of pgn and tPA-laden μwheels to the fibrin front and the local plasminogen and tPA concentrations (Eqns 5, 7, and 9). This model permits the prediction of plasminogen and tPA concentrations over time for a given number of mMSN/bead.

We used the model to calculate the percentage of time during a 60 min experiment in the AF zones defined in Fig. 2B as concentration ratios where lysis rates exceed 13.5 *μ*m/min. Coverage of 10-20 yielded the most favorable results with both species being maintained at target concentrations for at least 45 min (Fig. 5B). The model predicts that lower coverage results in faster plasminogen depletion than replenishment, while high coverage limits the available tPA on the bead surface.

To control the coverage, we varied the mixing time of the streptavidin beads and biotinylated mMSN (Fig. 5C). The coverage is 0-2 after 3 hr and 3-5 after 48 hr, a relatively slow association rate likely limited by steric hindrance at the bead surface. To achieve higher coverage, a 4 mT uniform magnetic field was applied across the sample, roughly doubling the coverage ratio for all mixing times (Fig. 5C). The distribution of coverage for a mixing time of 24 hr and a 4 mT field is shown in Fig. 5D. An average coverage of 8 mMSN/bead was achieved in 48 hr with a 4 mT field, a ratio where μwheels are predicted to achieve lysis in the AF zone for at least 40% of 1 hr experiment. This mixing time and magnetic field were used to synthesize pgn-tPA-beads for fibrinolysis experiments.

### Fibrinolysis with μwheel co-delivery of tPA and plasminogen

Fibrinolysis experiments were conducted on plasma clots with four formulations: free tPA, tPA beads, tPA-beads and pgn-mMSN (uncoupled), and pgn-tPA-beads (coupled) (Fig. 6A, Video S1). Particles with an initial activity equivalent to 5 nM tPA or free tPA (50 nM) were injected 1 mm from the fibrin front and a rotating magnetic field (6.2 mT, 10 Hz) was used to roll magnetic particles and their μwheel assemblies to the fibrin gel front. Free tPA diffused slowly to the fibrin front, resulting in negligible lysis for the first 20 min of the experiment (Fig. 6B). Our model estimates that the concentration of tPA at the fibrin front after 60 min is only 15 nM, indicating that transport is the rate-limiting process in this case (Fig. S1A). Once measurable, lysis proceeded at a linear rate of 0.42 ± 0.13 *μ*m/min for the duration of the experiment (Fig. 6B,C).

**Figure 6:**
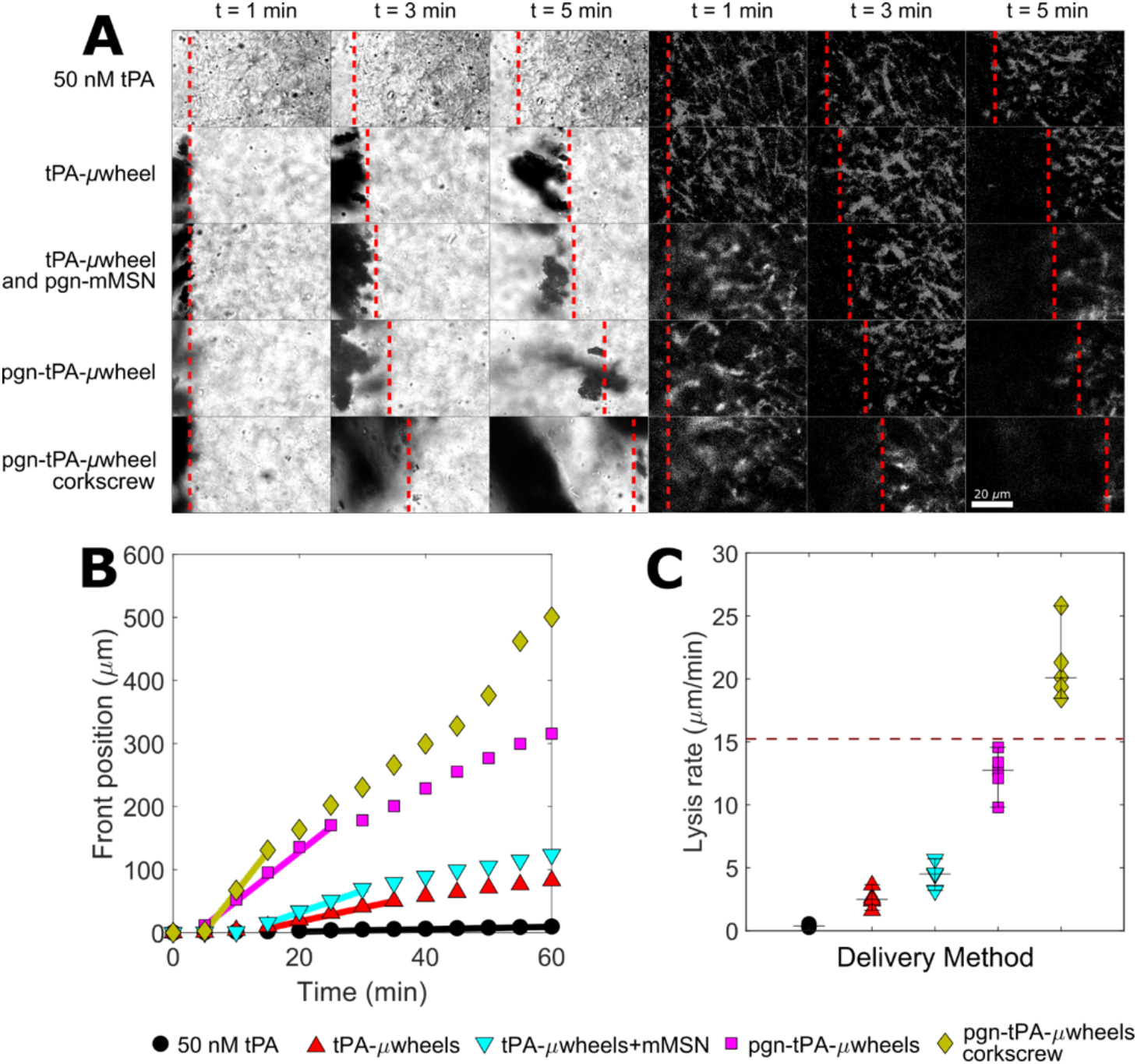
A) Time lapse of lysis of plasma clots using different μwheel formulations. Brightfield at left, and fluorescently labeled fibrin(ogen) at right. Dashed lines indicate the front position. B) Representative lysis curves for each formulation. Lines indicate periods of maximum lysis rates lasting for a minimum of 10 min. C) Maximum lysis rates for each formulation. Corkscrew condition uses pgn-tPA-*μ*wheels. Dashed line indicates biochemical speed limit (see Fig. 2B).

Experiments using tPA-beads were characterized by a shorter lag time (~10 min) than tPA as μwheels translate and accumulate at the fibrin front faster than diffusion (Fig. 6B). After this lag time, lysis rate accelerates as the accumulation of tPA-μwheels increases until it reaches a maximum of 2.9 ± 0.7 *μ*m/min (Fig. 6C) and then decreases after ~35 min. Our mathematical model estimates that tPA reaches 250 nM in 5 min and that plasminogen is depleted after 40 min (Fig. S1B) with tPA-μwheels. When tPA-μwheels are supplemented with uncoupled pgn mMSN, the maximum lysis rate increases to 4.5 ± 1.2 *μ*m/min (Fig. 6B). This relatively modest increase compared to tPA-beads alone is due to a limited amount of plasminogen replenishment; after 60 min, the plasminogen concentration is estimated to be 0.3 μM at the clot front (Fig. S1C).

The lysis rate of pgn-tPA-μwheels reaches a maximum rate of 13.6 ± 0.9 *μ*m/min (Fig. 6C) for the first 30 min of the experiment (Fig. 6B), consistent with the measured plasminogen release kinetics (Fig. 4C). The lag time is reduced for pgn-tPA-μwheels because of their increased average translation speed relative to tPA-μwheels (Fig. 4A-B). After 30 min, the pgn tPA-μwheel lysis rate approaches the rate of tPA-μwheels since most plasminogen has been released. Notably, this formulation reaches the biochemical speed limit of ~15 *μ*m/min (Fig. 2B) suggesting that both tPA and plasminogen are near their target concentrations. The mathematical model predicts that tPA reaches 250 nM after 10 min and plasminogen concentration remains above 1 μM for the duration of the experiment (Fig. S1D).

In order to break the biochemical speed limit, we used the magnetic field to manipulate *μ*wheels along a corkscrew trajectory as we have previously demonstrated that this allows μwheels to penetrate into clots.^13^ Pgn-tPA-μwheels following a corkscrew trajectory (Video S2) are 40-fold more effective fibrinolytics than 50 nM tPA and reach a maximum lysis rate of 20.3 ± 1.0 *μ*m/min (Fig. 6C). As μwheels burrow their way into a clot (Video S3), new fibrin fibers are available for binding of tPA and plasmin(ogen) that are not accessible during surface erosion. This mechanical penetration combined with high tPA and plasminogen concentration yields an inside-out lysis pattern that exceeds the maximum lysis rate achieved using only biochemical means (Fig. 2B).

## Discussion

The speed of fibrinolysis is governed by a series of rate-limiting transport and biochemical reaction steps. We have attempted to overcome each of those steps with the co delivery of a fibrinolytic enzyme, tPA, and its substrate, plasminogen, with a magnetically powered microbot. We measured the biochemical speed limit of fibrinolysis using different combinations of tPA and plasminogen concentrations and found that we can break that limit using this co-delivery strategy in combination with mechanical action created through corkscrew motion of μwheels.

The first rate-limiting step that must be overcome to achieve thrombolysis is the transport of fibrinolytics like tPA to a clot. Systemic delivery of fibrinolytic agents relies on diffusion to reach an impermeable occlusive clot,^24^ which is slow compared to active modes of transport or convection. We overcome this transport barrier by immobilizing tPA on magnetic beads which are assembled into μwheels that roll along a straight path to the clot. Using the active transport reduced the lag time to the start of fibrinolysis. Active transport also provides the opportunity to have a subtherapeutic circulating concentration of tPA immobilized on beads that accumulates at the clot at very high concentration.

The second rate-limiting step is the binding of tPA to fibrin. This step can be accelerated by increasing the concentration of tPA in systemic or intraarterial delivery. However, this approach is limited by the bleeding risk associated with high systemic concentrations of tPA. Using a μwheel delivery strategy offers an alternative solution because μwheels can be injected at subtherapeutic systemic concentration of tPA and subsequently localize at high concentrations at the clot interface. For example, in this study the tPA activity on our injected tPA-pgn-beads was comparable to 5 nM of free tPA, while the tPA activity of the accumulated μwheels at the clot interface is > 250 nM. tPA immobilized on microparticles is also less likely to enter the extravascular space due to their size and therefore has the potential to be less neurotoxic in stroke applications.

At sufficiently high tPA concentrations, plasminogen is the limiting factor in fibrinolysis speed.^49^ Once the fibrin front is replete with tPA, increasing tPA concentration further inhibits fibrinolysis because tPA both consumes the available plasminogen and competes with plasmin and plasminogen for fibrin binding sites. Here, we demonstrate that this limitation can not only be overcome by co-delivering a plasminogen payload with tPA using a microbot platform but can also approach the biochemical speed limit.

Both tPA and plasminogen have high affinity for fibrin fibers which limits their penetration into a clot leading to surface erosion of a clot with minimal penetration into low permeability clots. Thus, the last rate limiting step to overcome is the number of accessible fibrin binding sites.^55^ We have previously shown the potential to access binding sites within the interior of a clot, and a bulk erosion from the inside-out, using a μwheel corkscrew motion.^13^ Here we show that the corkscrew motion, in combination with co-delivery of tPA and plasminogen, breaks the biochemical speed limit by allowing μwheels to penetrate into the clot.

Using magnetically actuated pgn-tPA-μwheels, we have overcome each rate limiting step and achieved lysis rates that exceed even supratherapeutic concentrations of tPA. Our 40-fold increase in lysis rate over therapeutic concentrations of 50 nM tPA is a larger relative increase compared to other approaches that use magnetic particles as adjuvants or drug delivery vehicles.^42^ For example, using magnetic microparticles to augment tPA penetration into clots approximately doubles thrombolysis rates,^56^ a rate that can be further enhanced with the addition of ultrasound. Magnetic rod mixers can enhance the mass transfer of tPA such that fibrinolysis at lower concentrations of tPA can approach, but not exceed, that of supratherapeutic concentrations.^44^ However, strategies that increase the accessibility of local tPA ultimately run up against the limit of plasminogen availability. Because our approach uses beads encapsulating iron oxide, further enhancement of fibrinolysis rate could be achieved using high frequency magnetic fields to induce local hyperthermia, which has been shown to accelerate fibrinolysis in vitro and in vivo using tPA-functionalized iron oxide nanocubes.^45^ Taken together, the combination of multifunctional magnetic particles combined with dynamic magnetic fields has great potential to improve the rate of recanalization of occluded blood vessels, especially in cases like lacunar strokes where clots are not accessible to mechanical thrombectomy devices.

## Materials and Methods

### Materials

10 nm (II,III) iron oxide nanoparticles (CAS 900084), cetyltrimethylammonium bromide (CTAB, CAS 57-09-0), sodium dodecyl sulfate (SDS, CAS 151-21-3), sodium borohydride (NaBH4, CAS 16940-66-2), bovine serum albumin (BSA, CAS 9048-46-A), fibrinogen from human plasma (CAS 9001-32-5), and human plasminogen (SRP6518) were purchased from Sigma-Aldrich (St. Louis, MO). Ethyl acetate (142-89-2), ammonium hydroxide (1336-21-6) were obtained from Spectrum (New Brunswick, NJ). (Zeba desalting columns (89882), EZ Link™ Sulfo-NHS-Biotin kits (A39256), Dynabeads™ MyOne™ Streptavidin T1 (65601), tetraethyl orthosilicate (TEOS, O46174), and Alexa 555 (A20174) were obtained from Thermo Fisher Scientific (Waltham, MA). Normal pooled plasma (NPP) was purchased from George King Bio-Medical, Inc. (Overland Park, KS). Human alpha thrombin was obtained through Enzyme Research Laboratories (South Bend, IN). Recombinant human tPA was purchased from Abcam (Cambridge, UK).

### Synthesis of magnetic mesoporous silica nanoparticles (mMSN)

Synthesis of mMSN was adapted from Suteewong et al.^57^ where 15 mg of 10 nm iron oxide nanoparticles in chloroform were passivated in 54.8 mM CTAB via 5 min homogenization. The resulting emulsion was heated to 70°C for 10 min to evaporate the chloroform before being diluted 20x in 18.2 MΩ-cm DI water. Ethyl acetate, ammonium hydroxide, and TEOS at 0.062 M, 0.46 M and 0.015 M respectively were allowed to react for 8 min before being neutralized using 2 M HCl. The resulting particles were calcinated at 500 °C for 8 hr to remove the surfactant template before being resuspended in DI water with 0.1 wt% SDS. The entire reaction was carried out in a directional, no-gradient 4 mT magnetic field to bias the orientation of the iron oxide domains and maximize magnetic response. For some samples, mMSN were etched with NaBH_4_ for 4 hr to increase pore diameter after neutralization.

### Characterization of mMSN

Scanning transmission electron microscopy (STEM) images were captured using a Talos F200X microscope at an accelerating voltage of 200 keV. High-angle annular dark field energy dispersive X-ray (HAADF-EDS) spectra were also generated. Exposure times were greater than 30 min to allow for estimation of sample composition within a 90% confidence interval. Pore size distribution was measured using a Micromeritics Tristar 3000 sorptometer and calculated using the Barrett-Joyner-Halenda (BJH) method.^58^ Magnetization was characterized in an MPMS3 Quantum Design magnetometer at 25 °C from −0.5 to 0.5 T.

### Co-functionalization of Dynabeads™ with tPA and pgn-mMSN

To functionalize mMSN with primary amines, mMSN were incubated in 0.04 M 3-aminopropyltriethoxysilane (APS) in ethanol for 2 hr at room temperature, and then 1 hr at 80 °C.^59^ The mMSN were then washed with ethanol 4 times via centrifugation at 1500*g* for 1 min and finally resuspended in a 1:1 mixture of water and dimethyl sulfoxide (DMSO) (pH = 4.7). Then mMSN at 10^14^/*μ*L were mixed with 2 mL of 0.06M biotin in DMSO under sonication for 2 hr at room temperature. The resulting solution was washed 4 times with tris-buffer solution (pH = 7.4) via centrifugation at 1500*g* for 1 min. After the final wash cycle, the biotinylated mMSN were resuspended with 10^7^/*μ*L beads in TBS for 4-48 hr. A magnet was used to separate the beads from the unbound mMSN in four separate washing cycles. Each time, beads with covalently attached mMSN (studded beads) were resuspended in TBS. Biotinylated tPA (200 μg/mL) was added to the solution and the studded beads allowed to incubate for 12 hr at 2 °C before four more wash cycles. Finally, the studded beads were incubated in 10 μM plasminogen in TBS for 12 hr at 4°C. Immediately before use, the resulting beads co-functionalized with tPA and pgn-mMSN (pgn-tPA-beads) were washed three times in TBS to remove excess plasminogen.

### Magnetic field induced assembly and translation

Under the influence of a rotating magnetic field, tPA-beads and pgn-mMSN spontaneously assemble into μwheels.^53^ To direct the μwheels to the fibrin front, a rotating AC magnetic field was generated using five air-cored solenoids (51 mm inner diameter) with 336 turns each. An analog output card (National Instruments (NI), NI-9623, Austin, TX) controlled by Matlab, generated a current that was subsequently amplified (Behringer, EP2000, Willich, Germany). The final current passing through each coil, measured using an analog input card (NI-USB 6009), was 2 A, resulting in a field strength of 6.2 mT at the sample. The frequency of the rotating field was 10 Hz. For corkscrew experiments, the heading direction and camber angle for each magnetic field were shifted over time such that the beads followed a forward-biased spiral path with a 1s frequency. μWheel velocity was tracked using brightfield microscopy on a Prior Open Stand microscope (Prior Scientific, Cambridge, UK) with a PCO Panda camera (PCO Imaging, Kelheim, Germany).

### Loading and release of plasminogen

mMSN (10^6^/μL) were incubated in a 0.5 mL solution containing 10 μM plasminogen in PBS for 24 hr. Immediately before use, the pgn-mMSN were removed from solution via centrifugation at 5000*g* for 5 min, washed three times in 0.5 mL PBS, and resuspended in either PBS with 0.1 wt% SDS or plasma with 1 wt% BSA. A standard curve for plasminogen absorbance at 285 nm (A285) in PBS (0-20 μM) was measured with a UV-Vis spectrophotometer (Genesys 10S, ThermoFisher). Pgn-mMSN (10^5^/μL) were placed in 0.5 mL PBS. Every 5 min for 1 hr, the sample was centrifuged and the supernatant collected into a cuvette. Once the A285 was measured, the supernatant was used to resuspend the pgn-mMSN. A285 measurements were compared against a standard curve to calculate the release of plasminogen over time.

### Biotinylation of tPA

Recombinant tPA at 200 ug/mL was biotinylated with an EZ-Link Sulfo-NHS-Biotin kit and purified in Zeba Spin Desalting Columns according to manufacturer instructions. To prepare tPA functionalized beads (tPA-beads), 5 μL beads were incubated in 20 μL aliquots of 200 μg/mL biotinylated tPA at 4 °C for 12 hr (tPA-beads). Excess tPA was removed with four washes via centrifugation at 1500 g for 1 minute. tPA-beads were resuspended in TBS with 1 wt% BSA.

### Measurement of tPA activity on functionalized beads

tPA (0.25, 0.5, 0.75 and 1 μM) and tPA-beads (10^5^ and 10^6^/μL) were incubated with excess (100 μM) fluorogenic substrate for tPA (SN-18, Haematologic Technologies Inc., Essex Junction, VT). For solvated tPA experiments, fluorescent intensity was measured in a microplate reader (Biotek Synergy H1, BioTek U.S. Winooski, VT) every minute for 1 hr. For tPA-bead experiments, the solution was rotated on a carousel to prevent bead settling. In 5 min intervals, beads were extracted from solution using a permanent magnet and the fluorescent intensity of the supernatant measured. Beads and supernatant were remixed after each measurement to ensure experimental continuity.

### Modeling tPA and plasminogen concentrations at the lysis front

Diffusion of tPA was calculated using Fick’s Law for diffusion into a semi-infinite medium (Eqn 1):

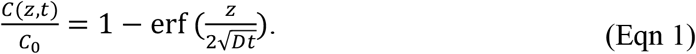

*C*(*z, t*) is the concentration at time *t* and location *z* of tPA, *C*_0_ is the injected concentration, and *D* is the diffusion constant (*D* = 5 * 10,^−11^ m^2^/s).^60^ To predict the concentrations of tPA and plasminogen over time for delivered species, we measured the velocity distribution of drug-laden pgn-tPA-μwheels as they approached the fibrin front. μWheels were driven in a single direction for 60 min under a 6 mT magnetic field at 10 Hz. The velocity of each individual bead was quantified using a macro for the Fiji distribution of ImageJ with the resulting distribution *f(v)* lognormal (Eqn 2) or Gaussian (Eqn 3):

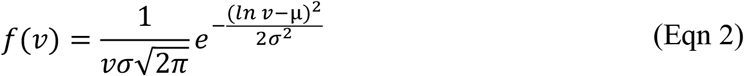

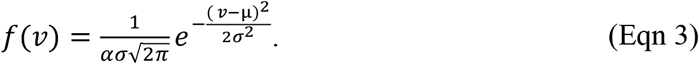

Here, μ is the arithmetic mean, *σ*^2^(the variance, and *α* a parameter that accounts for skew. Given these distribution functions, the probability that μwheels reach a point *z* at a given time *t* is:

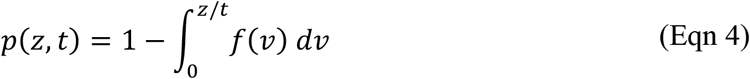

where *z* is the distance in microns between the fibrin front and bead injection site (1 mm). We define a rectangular control volume encompassing the area within *w =* 100 μm of the fibrin front. The concentration of beads that accumulate in that region is:

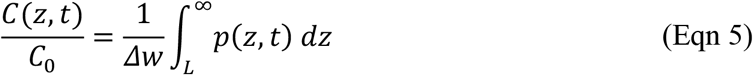

where *L* = 1000 μm is the distance separating the bead injection site from the fibrin front. Concentrations of tPA and plasminogen are calculated according to the following ordinary differential equations:

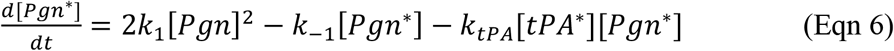

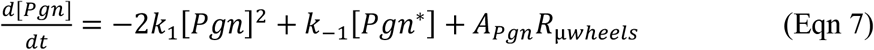

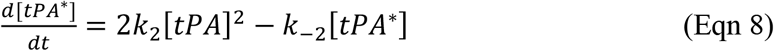

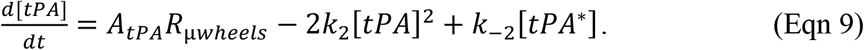

Here, [*Pgn*] is the concentration of free plasminogen, [*Pgn\**] is the concentration of plasminogen bound to fibrin, [*tPA*] is the concentration of free tPA and [*tPA\**] is the concentration of tPA bound to fibrin. The rate constants *k*_1_ and *k*_−1_ are the on and off rates for plasminogen binding to fibrin, and *k*_1_ and *k*_−1_, are the on and off rates for tPA binding to fibrin. *k_tPA_* is the rate at which tPA converts fibrin bound plasminogen to plasmin. Rate constants have been reported as *k*_1_ = 1.72 μM^−1^s^−1^, *k*_−1_ = 3.8 s^−1^ 61 *k*_−1_ = 0.011 μM^−1^s^−1^, *k*_−1_ = 0.0036 s^−1^ and *k_tPA_* = 0.01 μM^−1^s^−1^.^62,63^ *A_Pgn_* and *A_tPA_* are coefficients representing the active concentrations of plasminogen released from pgn-mMSN and tPA immobilized on tPA-μwheels respectively and derived from experimental measurements of plasminogen release and tPA activity as described above. *R*_μ*wheels*_ is the rate of accumulation of beads at the fibrin front and is the derivative with respect to time of Eqn 2. When solved numerically using the Matlab *ode45* solver, these equations can be used to estimate the tPA and plasminogen concentrations over the timescale of the fibrinolysis experiments.

### Fabrication of microfluidic devices

A template for the microfluidic channel was 3D printed using a Formlabs Form 3 printer using clear stereolithography resin (RS-F2-GPCL-04, Formlabs, Somerville, MA). The device has two inlets for the central channel (h = 50 μm) and two auxiliary inlets for the side chamber (h = 100 μm). A polydimethylsiloxane (PDMS) mold was made from the template. The PDMS was prepared using a 10:1 ratio of Sylgard 184 silicone elastomer base to curing agent (Dow Corning Corporation, Midland, MI). The mold was degassed for 1 hr in a vacuum chamber before being cured for 1 hr at 80 °C. Once cured, the mold was washed for 5 min in acetone and 5 min in ethanol in a sonicator before being air-dried. The mold and a glass slide were treated with oxygen plasma (0.2 torr) for 90 s and immediately bonded. The bonded assembly was annealed and then placed in a convection oven at 80 °C for 24 hr.

### Fibrinolysis experiments

The geometry for the device is shown in Fig. S2A. The two auxiliary inputs were plugged to prepare for injection of 30 μg/mL of Alexa 555 labeled fibrinogen, 20 mM CaCl2 and 9 nM thrombin to form a fibrin gel in the right channel. The device was enclosed in a Petri dish for 30 min with a moist Kimwipe to prevent evaporation and to allow full gelation. Next, the auxiliary chamber was filled with NPP and a suspension of tPA-beads and pgn-mMSN was injected 1 mm from the fibrin front. Lysis experiments were recorded using TRITC fluorescence and brightfield microscopy through a PCO Panda camera at 5 min intervals through a 40X objective (NA 0.95, Plan APO).

## Acknowledgments

The authors acknowledge support from the National Institutes of Health under grants R21AI138214 and R01NS102465. D.D. was supported by an American Heart Association Predoctoral Fellowship Award 18PRE34070076. Illustrations made with BioRender.com.

## Supplementary Information

Video S1: Brightfield (left) and fluorescence (right) time lapses of the dissolution of plasma clots using soluble tPA and various *μ*wheel populations. Video playback is accelerated 12X.

Video S2: Comparison of direct and corkscrew trajectories for a *μ*wheel monomer driven by a 10 Hz, 6.2 mT rotating magnetic field.

Video S3: Dissolution of plasma clots using pgn-tPA-*μ*wheels in either direct or corkscrew modes. Front positions are indicated with red lines. For the corkscrew mode, green circles track penetrating pgn-tPA-*μ*wheels.

Figure S1: Estimates from mathematical model of tPA and plasminogen concentrations for free tPA and different forumulations of microwheels.

Figure S2: Schematic of microfluidic device used for fibrinolysis experiments.

## References

(1) Han, K.; Shields, C. W.; Velev, O. D. Engineering of Self-Propelling Microbots and Microdevices Powered by Magnetic and Electric Fields. Adv. Funct. Mater. 2018, 28 (25), 1–14. https://doi.org/10.1002/adfm.201705953.

(2) Yang, T.; Tasci, T. O.; Neeves, K. B.; Wu, N.; Marr, D. W. M. Magnetic Microlassos for Reversible Cargo Capture, Transport, and Release. Langmuir 2017, 33 (23), 5932–5937. https://doi.org/10.1021/acs.langmuir.7b00357.

(3) Yang, T.; Sprinkle, B.; Guo, Y.; Qian, J.; Hua, D.; Donev, A.; Marr, D. W. M.; Wu, N. Reconfigurable Microbots Folded from Simple Colloidal Chains. Proc. Natl. Acad. Sci. U. S. A. 2020, 117 (31), 18186–18193. https://doi.org/10.1073/pnas.2007255117.

(4) Ma, F.; Wang, S.; Wu, D. T.; Wu, N. Electric-Field–Induced Assembly and Propulsion of Chiral Colloidal Clusters. Proc. Natl. Acad. Sci. 2015, 112 (20), 6307–6312. https://doi.org/10.1073/pnas.1502141112.

(5) Sen, A.; Ibele, M.; Hong, Y.; Velegol, D. Chemo and Phototactic Nano/Microbots. Faraday Discuss. 2009, 143, 9–14. https://doi.org/10.1039/b916271m.

(6) Aghakhani, A.; Yasa, O.; Wrede, P.; Sitti, M. Acoustically Powered Surface-Slipping Mobile Microrobots. Proc. Natl. Acad. Sci. U. S. A. 2020, 117 (7), 3469–3477. https://doi.org/10.1073/pnas.1920099117.

(7) Chen, X. Z.; Jang, B.; Ahmed, D.; Hu, C.; De Marco, C.; Hoop, M.; Mushtaq, F.; Nelson, B. J.; Pané, S. Small-Scale Machines Driven by External Power Sources. Adv. Mater. 2018, 30 (15), 1–22. https://doi.org/10.1002/adma.201705061.

(8) Challis, L. J. Mechanisms for Interaction between RF Fields and Biological Tissue. Bioelectromagnetics 2005, 26 (SUPPL. 7), 98–106. https://doi.org/10.1002/bem.20119.

(9) Martel, S. Presenting a New Paradigm in Cancer Therapy: Delivering Therapeutic Agents Using Navigable Microcarriers. IEEE Pulse 2014, 5 (3), 48–55. https://doi.org/10.1109/MPUL.2014.2309581.

(10) Schmidt, C. K.; Medina-Sánchez, M.; Edmondson, R. J.; Schmidt, O. G. Engineering Microrobots for Targeted Cancer Therapies from a Medical Perspective. Nat. Commun. 2020, 11 (1), 1–18. https://doi.org/10.1038/s41467-020-19322-7.

(11) Ullrich, F.; Bergeles, C.; Pokki, J.; Ergeneman, O.; Erni, S.; Chatzipirpiridis, G.; Pané, S.; Framme, C.; Nelson, B. J. Mobility Experiments with Microrobots for Minimally Invasive Intraocular Surgery. Investig. Ophthalmol. Vis. Sci. 2013, 54 (4), 2853–2863. https://doi.org/10.1167/iovs.13-11825.

(12) Li, J.; Li, X.; Luo, T.; Wang, R.; Liu, C.; Chen, S.; Li, D.; Yue, J.; Cheng, S. H.; Sun, D. Development of a Magnetic Microrobot for Carrying and Delivering Targeted Cells. Sci. Robot. 2018, 3 (19). https://doi.org/10.1126/scirobotics.aat8829.

(13) Tasci, T. O.; Disharoon, D.; Schoeman, R. M.; Rana, K.; Herson, P. S.; Marr, D. W. M.; Neeves, K. B. Enhanced Fibrinolysis with Magnetically Powered Colloidal Microwheels. Small 2017, 1–11. https://doi.org/10.1002/smll.201700954.

(14) Weisel, J. W. Structure of Fibrin: Impact on Clot Stability. J. Thromb. Haemost. 2007, 5 (SUPPL. 1), 116–124. https://doi.org/10.1111/j.1538-7836.2007.02504.x.

(15) Van de Werf, F.; Arnold, A. E. R. Intravenous Tissue Plasminogen Activator and Size of Infarct, Left Ventricular Function, and Survival in Acute Myocardial Infarction. Br. Med. J. 1988, 297 (6660), 1374–1379. https://doi.org/10.1136/bmj.297.6660.1374.

(16) Goldhaber, S. Z.; Meyerovitz, M. F.; Braunwald, E.; Green, D.; Vogelzang, R. L.; Citrin, P.; Heit, J.; Sobel, M.; Brownell Wheeler, H.; Plante, D.; Kim, H.; Hopkins, A.; Tufte, M.; Stump, D. Randomized Controlled Trial of Tissue Plasminogen Activator in Proximal Deep Venous Thrombosis. Am. J. Med. 1990, 88 (3), 235–240. https://doi.org/10.1016/0002-9343(90)90148-7.

(17) Levine, M.; Hirsh, J.; Weitz, J.; Cruickshank, M.; Neemeh, J.; Turpie, A. G.; Gent, M. A Randomized Trial of a Single Bolus Dosage Regimen of Recombinant Tissue Plasminogen Activator in Patients with Acute Pulmonary Embolism. Chest 1990, 98 (6), 1473–1479. https://doi.org/10.1378/chest.98.6.1473.

(18) Sandercock, P.; Wardlaw, J. M.; Lindley, R. I.; Dennis, M.; Cohen, G.; Murray, G.; Innes, K.; Venables, G.; Czlonkowska, A.; Kobayashi, A.; Ricci, S.; Murray, V.; Berge, E.; Slot, K. B.; Hankey, G. J.; Correia, M.; Peeters, A.; Matz, K.; Lyrer, P.; Gubitz, G.; Blackwell,; et al. The Benefits and Harms of Intravenous Thrombolysis with Recombinant Tissue Plasminogen Activator within 6 h of Acute Ischaemic Stroke (the Third International Stroke Trial [IST-3]): A Randomised Controlled Trial. Lancet 2012, 379 (9834), 2352– 2363. https://doi.org/10.1016/S0140-6736(12)60768-5.

(19) Emberson, J.; Lees, K. R.; Lyden, P.; Blackwell, L.; Albers, G.; Bluhmki, E.; Brott, T.; Cohen, G.; Davis, S.; Donnan, G.; Grotta, J.; Howard, G.; Kaste, M.; Koga, M.; Von Kummer, R.; Lansberg, M.; Lindley, R. I.; Murray, G.; Olivot, J. M.; Parsons, M.; Tilley, B.; Toni, D.; Toyoda, K.; Wahlgren, N.; Wardlaw, J.; Whiteley, W.; Del Zoppo, G. J.; Baigent, C.; Sandercock, P.; Hacke, W. Effect of Treatment Delay, Age, and Stroke Severity on the Effects of Intravenous Thrombolysis with Alteplase for Acute Ischaemic Stroke: A Meta-Analysis of Individual Patient Data from Randomised Trials. Lancet 2014, 384 (9958), 1929–1935. https://doi.org/10.1016/S0140-6736(14)60584-5.

(20) Byrne, R. M.; Taha, A. G.; Avgerinos, E.; Marone, L. K.; Makaroun, M. S.; Chaer, R. A. Contemporary Outcomes of Endovascular Interventions for Acute Limb Ischemia. J. Vasc. Surg. 2014, 59 (4), 988–995. https://doi.org/10.1016/j.jvs.2013.10.054.

(21) Hoylaerts, M.; Rijken, D. C.; Lijnen, H. R.; Collen, D. Kinetics of the Activation of Plasminogen by Human Tissue Plasminogen Activator. Role of Fibrin. J. Biol. Chem. 1982, 257 (6), 2912–2919. https://doi.org/10.1016/S0021-9258(19)81051-7.

(22) Francis, C. W.; Marder, V. J.; Barlow, G. H. Plasmic Degradation of Crosslinked Fibrin. Characterization of New Macromolecular Soluble Complexes and a Model of Their Structure. J. Clin. Invest. 1980, 66 (5), 1033–1043. https://doi.org/10.1172/JCI109931.

(23) Diamond, S. L. Engineering Design of Optimal Strategies for Blood Clot Dissolution. Annu. Rev. Biomed. Eng. 1999, 1 (1), 427–461. https://doi.org/10.1146/annurev.bioeng.1.1.427.

(24) Labiche, L. A.; Malkoff, M.; Alexandrov, A. V. Residual Flow Signals Predict Complete Recanalization in Stroke Patients Treated with TPA. J. Neuroimaging 2003, 13 (1), 28– 33. https://doi.org/10.1177/1051228402239714.

(25) Sabovic, M.; Blinc, A. Biochemical and Biophysical Conditions for Blood Clot Lysis. Pflugers Arch. Eur. J. Physiol. 2000, 440 (SUPPL. 5), 134–136. https://doi.org/10.1007/s004240000035.

(26) Chandler, W. L.; Alessi, M. C.; Aillaud, M. F.; Henderson, P.; Vague, P.; Juhan-Vague, I. Clearance of Tissue Plasminogen Activator (TPA) and TPA/Plasminogen Activator Inhibitor Type 1 (PAI-1) Complex. Circulation 1997, 96 (3), 761–768. https://doi.org/10.1161/01.cir.96.3.761.

(27) von Kummer, R. Early Major Ischemic Changes on Computed Tomography Should Preclude Use of Tissue Plasminogen Activator. Stroke 2003, 34 (3), 820–821. https://doi.org/10.1161/01.str.0000059430.55671.56.

(28) Henderson, S. J.; Weitz, J. I.; Kim, P. Y. Fibrinolysis: Strategies to Enhance the Treatment of Acute Ischemic Stroke. J. Thromb. Haemost. 2018, 16 (10), 1932–1940. https://doi.org/10.1111/jth.14215.

(29) Campbell, B. C. V.; Mitchell, P. J.; Churilov, L.; Yassi, N.; Kleinig, T. J.; Dowling, R. J.; Yan, B.; Bush, S. J.; Dewey, H. M.; Thijs, V.; Scroop, R.; Simpson, M.; Brooks, M.; Asadi, H.; Wu, T. Y.; Shah, D. G.; Wijeratne, T.; Ang, T.; Miteff, F.; Levi, C. R.; Davis, S. M.; et al. Tenecteplase versus Alteplase before Thrombectomy for Ischemic Stroke. N. Engl. J. Med. 2018, 378 (17), 1573–1582. https://doi.org/10.1056/nejmoa1716405.

(30) Logallo, N.; Novotny, V.; Assmus, J.; Kvistad, C. E.; Alteheld, L.; Rønning, O. M.; Thommessen, B.; Amthor, K. F.; Ihle-Hansen, H.; Kurz, M.; Tobro, H.; Kaur, K.; Stankiewicz, M.; Carlsson, M.; Morsund, Å.; Idicula, T.; Aamodt, A. H.; Lund, C.; Næss, H.; Waje-Andreassen, U.; Thomassen, L. Tenecteplase versus Alteplase for Management of Acute Ischaemic Stroke (NOR-TEST): A Phase 3, Randomised, Open-Label, Blinded Endpoint Trial. Lancet Neurol. 2017, 16 (10), 781–788. https://doi.org/10.1016/S1474-4422(17)30253-3.

(31) Sakharov, D. V.; Nagelkerkel, J. F.; Rijken, D. C. Rearrangements of the Fibrin Network and Spatial Distribution of Fibrinolytic Components during Plasma Clot Lysis: Study with Confocal Microscopy. J. Biol. Chem. 1996, 271 (4), 2133–2138. https://doi.org/10.1074/jbc.271.4.2133.

(32) Longstaff, C.; Thelwell, C.; Williams, S. C.; Silva, M. M. C. G.; Szabó, L.; Kolev, K. The Interplay between Tissue Plasminogen Activator Domains and Fibrin Structures in the Regulation of Fibrinolysis: Kinetic and Microscopic Studies. Blood 2011, 117 (2), 661– 668. https://doi.org/10.1182/blood-2010-06-290338.

(33) Donnan, G. A.; Davis, S. M.; Parsons, M. W.; Ma, H.; Dewey, H. M.; Howells, D. W. How to Make Better Use of Thrombolytic Therapy in Acute Ischemic Stroke. Nat. Rev. Neurol. 2011, 7 (7), 400–409. https://doi.org/10.1038/nrneurol.2011.89.

(34) Wang, Y. F.; Tsirka, S. E.; Strickland, S.; Stieg, P. E.; Soriano, S. G.; Lipton, S. A. Tissue Plasminogen Activator (TPA) Increase Neuronal Damage after Focal Cerebral Ischemia in Wild-Type and TPA-Deficient Mice. Nat. Med. 1998, 4 (2), 228–231. https://doi.org/10.1038/nm0298-228.

(35) Sakharov, D. V; Barrertt-Bergshoeff, M.; Hekkenberg, R. T.; Rijken, D. C. Fibrin Specificity of a Plasminogen Activator Affects the Efficiency of Fibrinolysis and Responsiveness to Ultrasound: Comparison of Nine Plasminogen Activators in Vitro. Thromb. Haemost. 1999, 81 (4), 605–612.

(36) Horrevoets, A. J. G.; Pannekoek, H.; Nesheim, M. E. A Steady-State Template Model That Describes the Kinetics of Fibrin Stimulated [Glu1] and [Lys78]Plasminogen Activation by Native Tissue Type Plasminogen Activator and Variants That Lack Either the Finger or Kringle-2 Domain. J. Biol. Chem. 1997, 272 (4), 2183–2191. https://doi.org/10.1074/jbc.272.4.2183.

(37) De Vries, C.; Veerman, H.; Koornneef, E.; Pannekoek, H. Tissue-Type Plasminogen Activator and Its Substrate Glu-Plasminogen Share Common Binding Sites in Limited Plasmin-Digested Fibrin. J. Biol. Chem. 1990, 265 (23), 13547–13552. https://doi.org/10.1016/s0021-9258(18)77382-1.

(38) Jung He Wu; Diamond, S. L. Tissue Plasminogen Activator (TPA) Inhibits Plasmin Degradation of Fibrin: A Mechanism That Slows TPA-Mediated Fibrinolysis but Does Not Require Α2 Antiplasmin or Leakage of Intrinsic Plasminogen. J. Clin. Invest. 1995, 95 (6), 2483–2490. https://doi.org/10.1172/jci117949.

(39) Kim, P. Y.; Tieu, L. D.; Stafford, A. R.; Fredenburgh, J. C.; Weitz, J. I. A High Affinity Interaction of Plasminogen with Fibrin Is Not Essential for Efficient Activation by Tissue Type Plasminogen Activator. J. Biol. Chem. 2012, 287 (7), 4652–4661. https://doi.org/10.1074/jbc.M111.317719.

(40) Onundarson, P. T.; Francis, C. W.; Marder, V. J. Depletion of Plasminogen in Vitro or during Thrombolytic Therapy Limits Fibrinolytic Potential. J. Lab. Clin. Med. 1992, 120 (1), 120–128.

(41) Baeten, K. M.; Richard, M. C.; Kanse, S. M.; Mutch, N. J.; Degen, J. L.; Booth, N. A. Activation of Single-Chain Urokinase-Type Plasminogen Activator by Platelet-Associated Plasminogen: A Mechanism for Stimulation of Fibrinolysis by Platelets. J. Thromb. Haemost. 2010, 8 (6), 1313–1322. https://doi.org/10.1111/j.1538-7836.2010.03813.x.

(42) Disharoon, D.; Marr, D. W. M.; Neeves, K. B. Engineered Microparticles and Nanoparticles for Fibrinolysis. J. Thromb. Haemost. 2019, 17 (12), 2004–2015. https://doi.org/10.1111/jth.14637.

(43) Hu, J.; Huang, W.; Huang, S.; ZhuGe, Q.; Jin, K.; Zhao, Y. Magnetically Active Fe3O4 Nanorods Loaded with Tissue Plasminogen Activator for Enhanced Thrombolysis. Nano Res. 2016, 9 (9), 2652–2661. https://doi.org/10.1007/s12274-016-1152-4.

(44) Huang, L.; ZhuGe, Q.; Cheng, R.; Zhao, Y.; Mao, L.; Yang, B.; Jin, K.; Huang, W. Acceleration of Tissue Plasminogen Activator-Mediated Thrombolysis by Magnetically Powered Nanomotors. ACS Nano 2014, 8 (8), 7746–7754. https://doi.org/10.1021/nn5029955.

(45) Voros, E.; Cho, M.; Ramirez, M.; Palange, A. L.; De Rosa, E.; Key, J.; Garami, Z.; Lumsden, A. B.; Decuzzi, P. TPA Immobilization on Iron Oxide Nanocubes and Localized Magnetic Hyperthermia Accelerate Blood Clot Lysis. Adv. Funct. Mater. 2015, 25 (11), 1709–1718. https://doi.org/10.1002/adfm.201404354.

(46) Chen, J. P.; Yang, P. C.; Ma, Y. H.; Wu, T. Characterization of Chitosan Magnetic Nanoparticles for in Situ Delivery of Tissue Plasminogen Activator. Carbohydr. Polym. 2011, 84 (1), 364–372. https://doi.org/10.1016/j.carbpol.2010.11.052.

(47) Ma, Y. H.; Wu, S. Y.; Wu, T.; Chang, Y. J.; Hua, M. Y.; Chen, J. P. Magnetically Targeted Thrombolysis with Recombinant Tissue Plasminogen Activator Bound to Polyacrylic Acid-Coated Nanoparticles. Biomaterials 2009, 30 (19), 3343–3351. https://doi.org/10.1016/j.biomaterials.2009.02.034.

(48) Yang, H. W.; Hua, M. Y.; Lin, K. J.; Wey, S. P.; Tsai, R. Y.; Wu, S. Y.; Lu, Y. C.; Liu, H. L.; Wu, T.; Ma, Y. H. Bioconjugation of Recombinant Tissue Plasminogen Activator to Magnetic Nanocarriers for Targeted Thrombolysis. Int. J. Nanomedicine 2012, 7, 5159– 5173. https://doi.org/10.2147/IJN.S32939.

(49) Rijken, D. C.; Sakharov, D. V. Basic Principles in Thrombolysis: Regulatory Role of Plasminogen. Thromb. Res. 2001, 103 (SUPPL. 1), 41–49. https://doi.org/10.1016/S0049-3848(01)00296-1.

(50) Deodhar, G. V.; Adams, M. L.; Joardar, S.; Joglekar, M.; Davidson, M.; Smith, W. C.; Mettler, M.; Toler, S. A.; Davies, F. K.; Williams, S. K. R.; Trewyn, B. G. Conserved Activity of Reassociated Homotetrameric Protein Subunits Released from Mesoporous Silica Nanoparticles. Langmuir 2018, 34 (1), 228–233. https://doi.org/10.1021/acs.langmuir.7b03310.

(51) Martin-Ortigosa, S.; Peterson, D. J.; Valenstein, J. S.; Lin, V. S. Y.; Trewyn, B. G.; Alexander Lyznik, L.; Wang, K. Mesoporous Silica Nanoparticle-Mediated Intracellular Cre Protein Delivery for Maize Genome Editing via LoxP Site Excision. Plant Physiol. 2014, 164 (2), 537–547. https://doi.org/10.1104/pp.113.233650.

(52) Martin-Ortigosa, S.; Valenstein, J. S.; Lin, V. S. Y.; Trewyn, B. G.; Wang, K. Gold Functionalized Mesoporous Silica Nanoparticle Mediated Protein and DNA Codelivery to Plant Cells via the Biolistic Method. Adv. Funct. Mater. 2012, 22 (17), 3576–3582. https://doi.org/10.1002/adfm.201200359.

(53) Tasci, T. O.; Herson, P. S.; Neeves, K. B.; Marr, D. W. M. Surface-Enabled Propulsion and Control of Colloidal Microwheels. Nat. Commun. 2016, 7, 10225. https://doi.org/10.1038/ncomms10225.

(54) Disharoon, D.; Neeves, K. B.; Marr, D. W. M. Ac/Dc Magnetic Fields for Enhanced Translation of Colloidal Microwheels. Langmuir 2019, 35 (9), 3455–3460. https://doi.org/10.1021/acs.langmuir.8b04084.

(55) Bannish, B. E.; Keener, J. P.; Fogelson, A. L. Modelling Fibrinolysis: A 3D Stochastic Multiscale Model. Math. Med. Biol. 2014, 31 (1), 17–44. https://doi.org/10.1093/imammb/dqs029.

(56) Torno, M. D.; Kaminski, M. D.; Xie, Y.; Meyers, R. E.; Mertz, C. J.; Liu, X.; O’Brien, W. D.; Rosengart, A. J. Improvement of in Vitro Thrombolysis Employing Magnetically Guided Microspheres. Thromb. Res. 2008, 121 (6), 799–811. https://doi.org/10.1016/j.thromres.2007.08.017.

(57) Suteewong, T.; Sai, H.; Lee, J.; Bradbury, M.; Hyeon, T.; Gruner, S. M.; Wiesner, U. Ordered Mesoporous Silica Nanoparticles with and without Embedded Iron Oxide Nanoparticles: Structure Evolution during Synthesis. J. Mater. Chem. 2010, 20 (36), 7807–7814. https://doi.org/10.1039/c0jm01002b.

(58) Barrett, E. P.; Joyner, L. G.; Halenda, P. P. The Determination of Pore Volume and Area Distributions in Porous Substances. I. Computations from Nitrogen Isotherms. J. Am. Chem. Soc. 1951, 73 (2), 373–380.

(59) Corricelli, M.; Depalo, N.; Di Carlo, E.; Fanizza, E.; Laquintana, V.; Denora, N.; Agostiano, A.; Striccoli, M.; Curri, M. L. Biotin-Decorated Silica Coated PbS Nanocrystals Emitting in the Second Biological near Infrared Window for Bioimaging. R. Soc. Chem. 2013, 00 (1–3), 1–10. https://doi.org/10.1039/x0xx00000x.

(60) Weigandt, K. M.; White, N.; Chung, D.; Ellingson, E.; Wang, Y.; Fu, X.; Pozzo, D. C. Fibrin Clot Structure and Mechanics Associated with Specific Oxidation of Methionine Residues in Fibrinogen. Biophys. J. 2012, 103, 2399–2407.

(61) Wiman, B.; Collen, D. On the Mechanism of the Reaction between Human Α2 Antiplasmin and Plasmin. J. Biol. Chem. 1979, 84 (18), 573–578.

(62) Tiefenbrunn, A. J.; Graor, R. A.; Robison, A. K.; Lucas, F. V.; Hotchkiss, A.; Sobel, B. E. Pharmacodynamics of Tissue-Type Plasminogen Activator Characterized by Computer Assisted Simulation. Circulation 1986, 73 (6), 1291–1299. https://doi.org/10.1161/01.CIR.73.6.1291.

(63) Bannish, B. E.; Chernysh, I. N.; Keener, J. P.; Fogelson, A. L.; Weisel, J. W. Molecular and Physical Mechanisms of Fibrinolysis and Thrombolysis from Mathematical Modeling and Experiments. Sci. Rep. 2017, 7 (1), 1–11. https://doi.org/10.1038/s41598-017-06383-w.

