## Supporting Information for "Breaking the fibrinolytic speed limit with microwheel co-delivery of tissue plasminogen activator and plasminogen"

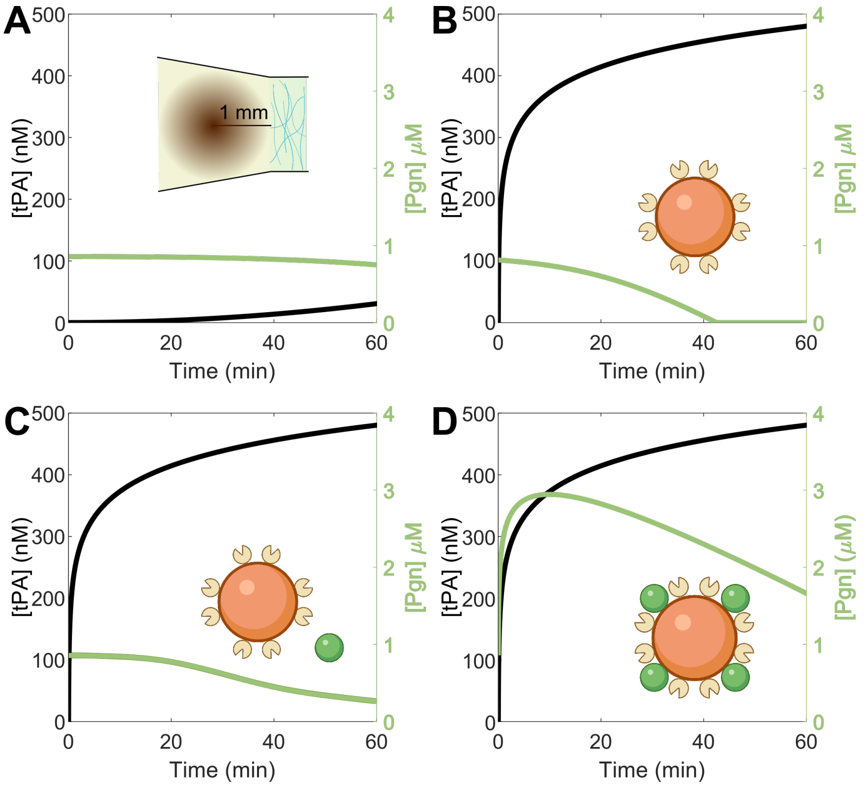


Figure S1: Estimates from mathematical model of tPA and plasminogen concentrations for free tPA and different forumulations of microwheels. A) Local concentration of tPA and plasminogen 1 mm from the injection of a 50 nM bolus of tPA and 1 μM plasminogen. B) Local concentration of tPA and plasminogen 1mm from the injection of $10^5$/μL tPA-beads. C) Local concentration of tPA and plasminogen 1mm from the injection of $10^5$/μL tPA-beads and $10^6$/μL pgn-mMSN. D) Local concentration of tPA and plasminogen 1 mm from the injection of $10^5$/μLpgn-tPA-beads.

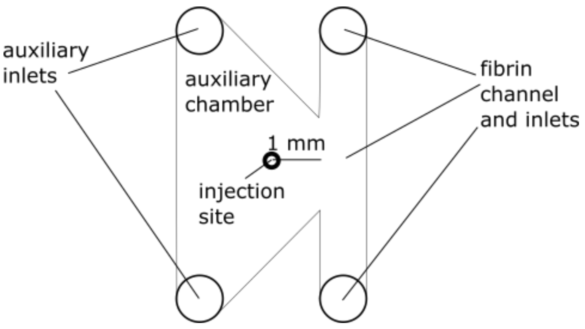


Figure S2: Schematic of microfluidic device used for fibrinolysis experiments.
